# Relapse-founding cancer persister cells in follicular lymphoma

**DOI:** 10.64898/2026.06.23.734001

**Authors:** Oscar Atkins, Miu Shing Hung, Ok-Ryul Song, Baizhi Chen, Bernard Maybury, Carina Edmondson, Bruno Tesson, Sarah Huet, Gilles Salles, Michael Howell, Hans Christian Reinhardt, Jude Fitzgibbon, Jessica Okosun, Lingling Zhang, Dinis Pedro Calado

## Abstract

Follicular lymphoma (FL) is an incurable, prototypical relapse-remitting cancer, implying the existence of therapy-persistent cells that survive frontline treatment and seed disease recurrence1-3. However, these persister cells remain difficult to study directly in patients because immediate post-treatment sampling is ethically and practically challenging. Using a genetically defined mouse model that allows sampling of persistent cells immediately after frontline R-CHOP therapy, we prospectively isolate and functionally define relapse-founding cancer persister cells (CPC). The CPC is an IgM⁺ memory-like B-cell with high germinal center re-entry capacity. This state represents a discrete component of a heterogeneous residual pool indicating that residual disease is polytypic and that relapse potential may depend on which cells persist rather than on residual tumour burden alone. By integrating mouse CPC with human FL datasets, we show that an analogous transcriptional programme is detectable at diagnosis and is enriched in patients with inferior clinical outcome across independent cohorts4,5. These findings support the concept that relapse risk is linked to a conserved, genotype-agnostic CPC programme present before therapy. To explore therapeutic vulnerabilities, we developed a scalable in-vitro platform that models the CPC-like state and used it to identify sensitivity to histone deacetylase inhibition. Romidepsin and panobinostat killed CPC-like cells in-vitro, and decreased therapy-persistent cells after R-CHOP treatment in-vivo and in patient-derived organoids. Together, these data define a tractable CPC state in FL, with a validated clinical readout and an immediately testable therapeutic entry point, opening CPC-directed strategies for durable FL control.

## Main

Follicular lymphoma (FL), the most common indolent non-Hodgkin lymphoma, is a prototypical relapsing-remitting cancer that remains incurable. Frontline rituximab-based immunochemotherapy (R-CHOP: Rituximab, Cyclophosphamide, Doxorubicin, Vincristine, Prednisolone; or R-bendamustine) induces remission in most patients. However, early relapse within 24 months (POD24) and transformation to aggressive diffuse large B-cell lymphoma are associated with poor outcomes. Because existing clinical^6,7^ and molecular tools^5,8,9^ have modest prospective utility, these patients can only be identified retrospectively^10,11^, limiting opportunities for risk-adapted early intervention. A cellular biomarker that is measurable at diagnosis and mechanistically linked to relapse would enable rational stratification.

Relapsing-remitting cancers imply a therapy-persistent population. In FL, longitudinal phylogenetics supports a relapse-founding cancer persister cell (CPC)^1–3^ that carries the hallmark t(14;18) translocation and early alterations in chromatin-modifying genes e.g., CREBBP and KMT2D^1–3,12^. CPCs can persist for years, including in otherwise healthy individuals^13–15^. Nonetheless, the CPC has never been isolated or functionally defined, precluding CPC-directed therapies^16^. A central obstacle is that immediate post-R-CHOP sampling of therapy-persistent cells is not feasible in patients.

Here we overcame this barrier with a mouse-first design that permitted immediate post-therapy sampling of therapy-persistent cells, enabling prospective isolation and functional definition of the relapse-founding CPC from a heterogeneous residual pool, thus separating “residual disease” from “relapse potential”. The CPC was an IgM⁺ memory B-cell (MBC)-like subset with high germinal center (GC) re-entry potential, consistent with GC-centred biology of FL pathogenesis^17–20^. We translated the CPC program to patients as a gene signature that mapped this state at diagnosis, stratified risk across independent cohorts^4,5^, suggesting a genotype-agnostic founder program with immediate utility for baseline profiling and longitudinal monitoring. Using a scalable CPC-like platform, we revealed dependencies on histone-deacetylase activity and demonstrated CPC killing using romidepsin and panobinostat, defining a tractable therapeutic axis for durable disease control.

### Frontline immunochemotherapy reveals persistent cells

Across integrated FL patient datasets (n=76), clonal alterations in histone-modifying genes were frequent and shared at diagnosis, relapse and transformation, placing them early in FL evolution^1–3,12^. CREBBP and KMT2D alterations, present in 67% and 62% of cases, respectively, were inferred to reside in the CPC reservoir, and 51% of patients harboured ≥2 chromatin-gene mutations, most commonly the CREBBP-KMT2D pair (**Fig. 1a,b**). These patterns implicate epigenetic dysregulation as a defining feature of the relapse-founder state.

**Figure 1:**
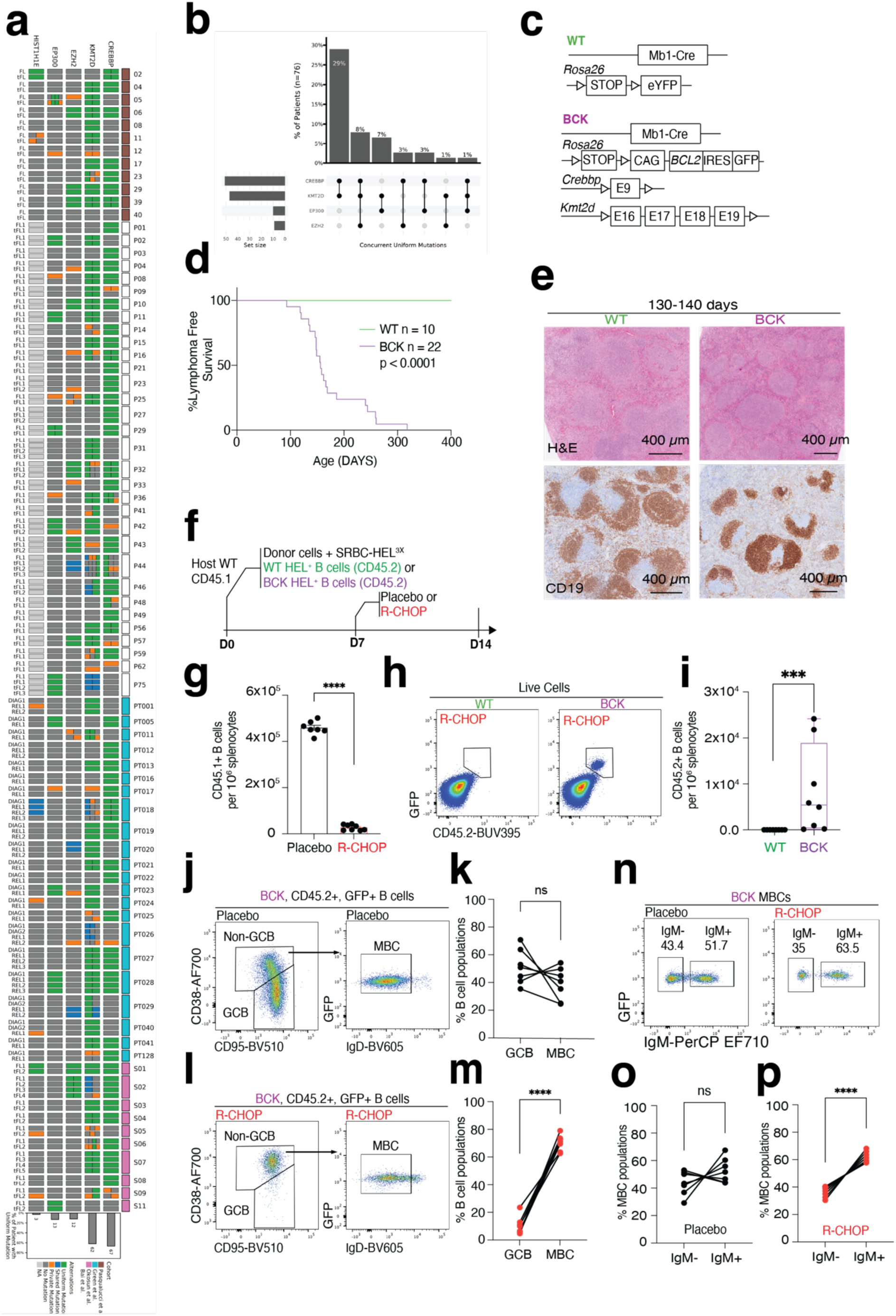
Frontline therapy reveals persistent cells. (a) Heatmap integrating publicly available exome sequencing data from paired diagnosis, relapsed FL, and transformed FL samples. **(b)** UpSet plot showing the frequency and co-occurrence of mutations in patients with ≥2 mutations in chromatin-modifying genes (*CREBBP*, *KMT2D*, *EP300*, and *EZH2*). **(c)** Schematic of allele configurations in WT and BCK mice. Both carry a Rosa26 allele with a *loxP-stop-loxP-eYFP* (WT) or *BCL2-IRES-GFP* (BCK) cassette. BCK mice additionally harbor loxP-flanked exon 9 of *CREBBP* and exons 16–19 of *KMT2D*. **(d)** Kaplan-Meier survival curves comparing BCK (n=22) and WT (n=10) mice. **(e)** Representative immunohistochemistry (IHC) images of spleen sections; scale bar = 400 μm. **(f)** Experimental timeline. **(g)** Absolute number of host-derived (CD45.1⁺) B-cells per 10⁶ splenocytes in mice treated with placebo (isotype control + saline) or R-CHOP. Mean ± SEM; Placebo n=7, R-CHOP n=8. **(h)** Flow cytometry gating strategy for donor-derived (CD45.2⁺) B-cells. **(i)** Absolute number of donor-derived (CD45.2⁺) B-cells per 10⁶ splenocytes in mice treated with R-CHOP. Mean ± SEM; WT n=8, BCK n=8. **(j, k)** Gating strategy and quantification of GC B-cells (GCB) and memory B-cells (MBC) among BCK donor cells in placebo-treated recipients. Mean ± SEM; n=7. **(l, m)** Gating strategy and quantification of GCB and MBC among BCK donor cells in R-CHOP–treated recipients. Mean ± SEM; n=8. **(n-p)** Gating strategy and quantification of IgM⁺ and IgM^neg^ BCK MBCs in placebo-and R-CHOP–treated mice. Mean ± SEM; placebo n=7, R-CHOP n=8. Each dot represents an individual mouse. Data in (d) two-tailed log-rank test, (g, k, m, o and p) are representative of two independent experiments. \**P* ≤ 0.05; \*\**P* ≤ 0.01; \*\*\**P* ≤ 0.001; \*\*\*\**P* ≤ 0.0001; ns, not significant (two-tailed Student’s *t*-test). (i) two-tailed Mann Whitney test. \*\*\**P* ≤ 0.001.

To model this state, we generated BCK mice (*Cd79a*-cre; *R26-BCL2^LSL^*; *Crebbp*^+/fl^; *Kmt2d*^fl/fl^)^21–24^, marking cre-loxP recombination via GFP expression (**Fig. 1c**). Unlike prior strategies^25^, this design drives BCL2 from the early pro-B stage, within the developmental window of t(14;18)^26^, and recapitulates the common combination of heterozygous *CREBBP* loss with homozygous *KMT2D* loss seen in FL^27,28^. Compared with *Cd79a*-cre; *R26-YFP^LSL^* controls (WT), BCK mice showed reduced survival (∼22 weeks), splenomegaly and FL-like architecture with hypercellular follicles (**Fig. 1d,e**).

Because FL pathogenesis is GC-centred^18^, we coupled BCK to the *SWHEL* Ig knock-in to synchronize and precisely control GC entry using antigen-specific hen-egg lysozyme (HEL) immunization^29,30^. We then performed competitive adoptive transfer of naïve BCK and WT CD45.2^+^ B-cells into fully immunocompetent, syngeneic CD45.1^+^ hosts (**Supp. Fig. 1a**); seeding donors at ≤2% of total B-cells to mirror the low mutant-cell frequency observed in otherwise healthy individual^13–15^. This setup enables unambiguous fate-mapping of donor clones while preventing confounding input from newly generated mutants. After HEL^29,30^ immunization, BCK cells drove hyperplastic GCs with reduced differentiation to MBCs and plasma cells and an increased dark-zone (DZ): light-zone (LZ) ratio (**Supp. Fig. 1b-g**), consistent with known roles of CREBBP and KMT2D in GC exit and B-cell differentiation^16,31–33^. Proliferation and apoptosis were both reduced relative to WT, aligning with BCL2’s anti-apoptotic and anti-proliferative activities^34^ (**Supp. Fig. 1i,k**).

To test whether we could identify therapy-persistent cells^1–3,12–15^, we transferred naïve BCK or WT B-cells from tumor-free donors (∼8 weeks old) into immunocompetent hosts. Seven days after immunization, mice received mouse-adapted R-CHOP; spleens were analysed on day 14 (**Fig. 1f**). R-CHOP efficiently depleted endogenous CD45.1⁺ B-cells and eradicated WT donor B-cells (**Fig. 1g-i**). In contrast a subset of BCK donor cells persisted (**Fig. 1h,i**) and were predominantly MBC-like (CD38^hi^ CD95^lo^ IgD^neg^), as GC BCK cells were efficiently cleared (**Fig. 1j-m**). The surviving cells showed pronounced IgM retention^19,35,36^ (**Fig. 1n-p**). These data implicate IgM expression in CPC persistence.

Together, the genetic, modelling and therapeutic-challenge experiments demonstrate that B-cells bearing CREBBP/KMT2D loss with BCL2 overexpression can survive frontline immunochemotherapy as an IgM⁺ MBC-like population, establishing a therapy-persistent state consistent with a relapse-founder identity.

### Residual disease resolves into three MBC-like states

To define the cells that persist after therapy, we profiled CD45.2⁺ donor BCK cells by scRNA-seq from R-CHOP-treated recipients, alongside BCK cells from placebo-treated mice and spontaneous BCK tumors (**Fig. 2a**). InferCNV analysis showed that R-CHOP largely removed cells with large-scale chromosomal alterations present in tumors, leaving the residual compartment comparatively copy-number-quiet (**Fig. 2b,c**). Thus, therapy-persistent cells are molecularly distinct from fully transformed cells, consistent with a progenitor-like state.

**Figure 2:**
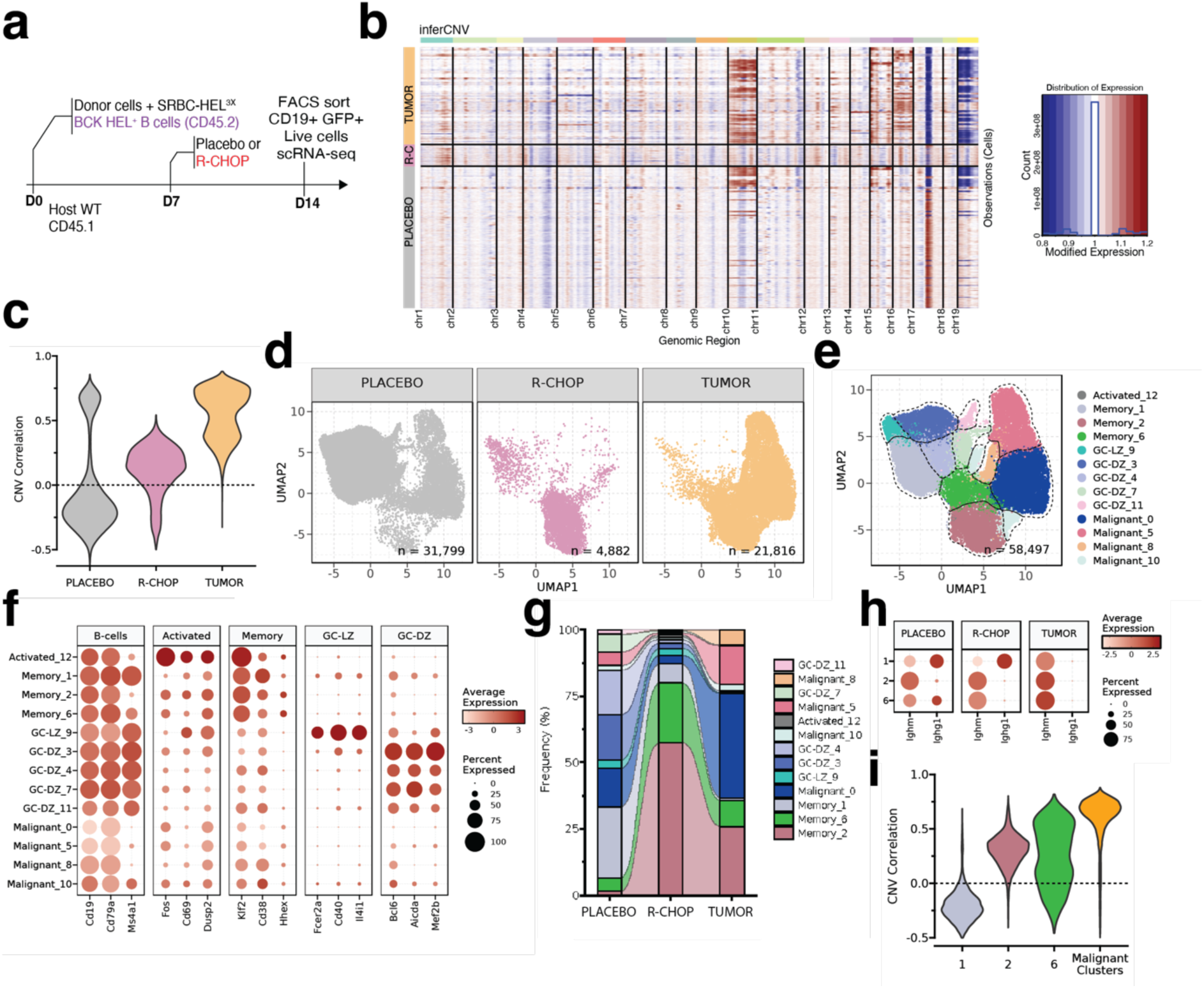
Residual disease resolves into three MBC-like states. (a) Experimental design. Naïve B-cells were isolated from the spleens of young BCK mice and transferred into CD45.1⁺ wild-type recipients. Recipients were immunized with SRBC-HEL3X and treated with R-CHOP or placebo on day 7. On day 14, mice were sacrificed, and donor-derived B-cells were FACS-sorted from spleens for single-cell RNA sequencing (scRNA-seq). In parallel, scRNA-seq was also performed on malignant B-cells isolated from aged BCK mice that had spontaneously developed overt FL. **(b)** inferCNV analysis across all cells, grouped by sample type. R-CHOP: CD45.2⁺ cells isolated from CD45.1⁺ recipient mice (n=9) following R-CHOP therapy; Placebo: BCK cells from placebo-treated mice (n=3), and Tumor: spontaneous lymphomas from untreated BCK mice (n=3). **(c)** Correlation between each cell’s CNV profile and the average CNV profile of tumor cells. **(d)** UMAP plot of integrated scRNA-seq data from all 58,497 cells, revealing 13 distinct transcriptional clusters. **(e)** Sample-type separation of cells: placebo-treated (BCK cells from placebo recipients), R-CHOP–persistent (BCK cells from R-CHOP–treated recipients), and tumor-derived (BCK tumor cells from aged mice). **(f)** Dot plot showing average gene expression and percentage of cells expressing selected marker genes across defined B-cell subtypes. **(g)** Cluster composition of each sample type, showing the proportion of cells belonging to each of the 13 identified clusters. **(h)** BCR isotype usage within memory clusters 1, 2, and 6. **(i)** CNV correlation analysis comparing the inferCNV profiles of individual cells in memory clusters 1, 2, and 6, and in malignant clusters 0, 5, 8, and 10, against the mean CNV profile of the tumor.

Joint integration of all datasets yielded a shared UMAP with 13 clusters (**Fig. 2d-f**). BCK cells from placebo-treated mice were dominated by DZ GC phenotypes, whereas ≥80% of residual BCK cells after R-CHOP populated three memory-like clusters (1, 2, 6) (**Fig. 2g**). Notably, cluster-2 retained IgM under both placebo and R-CHOP conditions (**Fig. 2h**), nominating an IgM-expressing MBC-like subset as a preferential survivor state. The same MBC-like clusters were detectable, at lower frequency, within spontaneous tumors, with cluster-2 again most represented (**Fig. 2g**). Copy-number patterns stratified these states: cluster-1 was largely genomically normal, whereas clusters-2 and-6 showed intermediate CNV burdens, consistent with early chromosomal change during pre-malignant progression (**Fig. 2i**).

Together, these data show that the post-R-CHOP reservoir is transcriptionally heterogeneous, memory-like, and largely copy-number–quiet, with an IgM-retaining memory subset emerging as the dominant therapy-persistent state.

### A conserved CPC program identifies high-risk patients

The presence of MBC-like therapy-persistent cells in mice prompted us to ask whether an analogous population exists in human FL. We analyzed a diagnostic FL scRNA-seq dataset with outcome data^37^ and mapped patient B-cells onto the mouse reference (**Fig. 3a**). Integrated alignment identified human counterparts of the BCK-derived states (**Fig.2e, Fig. 3a**), including the three MBC-like clusters (1, 2, 6) that are enriched after R-CHOP. Cluster-2, which uniquely retained IgM, was over-represented in tumors from patients who experienced POD24 (**Fig. 3b**), nominating a conserved IgM-retaining MBC-like persister state as a candidate CPC reservoir.

**Figure 3:**
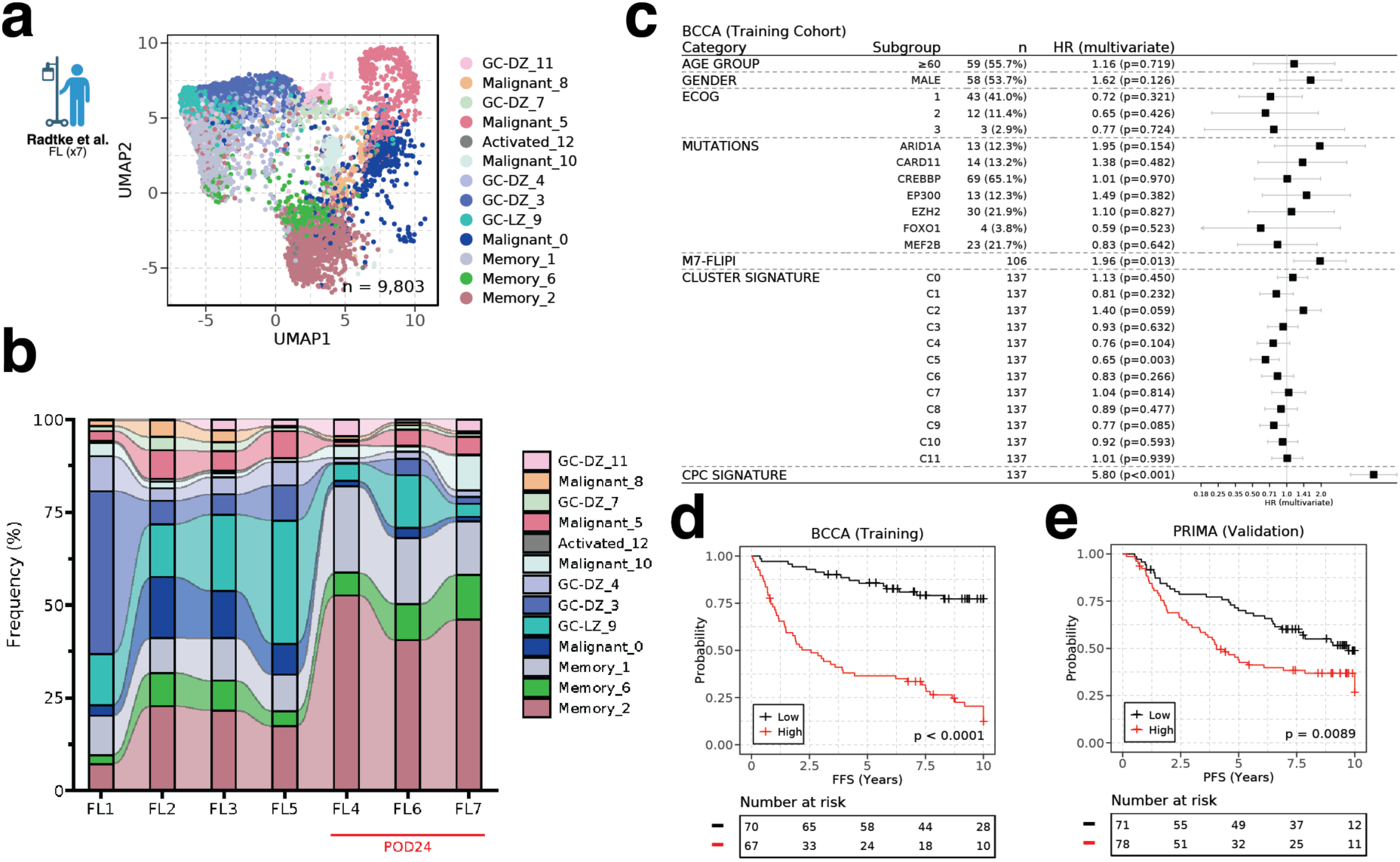
A conserved CPC program identifies high-risk patients. (a) Cross-species reference mapping of human FL scRNA-seq data from Radtke et a^37^. to mouse-defined clusters identified in our study. **(b)** Patient-level distribution of tumor cells across mouse-defined clusters in the Radtke dataset, showing inter-patient heterogeneity in cluster representation. **(c)** Prognostic analysis in the BCCA cohort: hazard ratios (HRs) for failure-free survival (FFS) associated with enrichment of differentially expressed genes from each of the 13 mouse-derived clusters. The HR for the optimized CPC signature derived from cluster-2 MBCs is shown in the bottom row. HR and p values calculated based on multivariate cox regression model (CoxPH). **(d, e)** Kaplan-Meier survival analyses of FFS (BCCA training cohort) and progression-free survival (PFS; PRIMA validation cohort) based on CPC signature enrichment. P value was calculated based on log likelihood ratio test.

To test prognostic relevance, we turned to RNA-seq from the BCCA^4^ trial cohort (n=137). Enrichment of a cluster-2 gene set associated with inferior failure-free survival (FFS; HR 1.40, p=0.059), whereas no other cluster program predicted adverse outcomes (**Fig. 3c**). We then used elastic-net regression to identify the subset of cluster-2 program genes most strongly linked to poor prognosis, generating a CPC signature (**Fig. 3c**). When applied to the BCCA cohort, the CPC signature stratified patients into distinct risk groups. High CPC signature enrichment (n=67/137, 48.9%) was associated with significantly worse 5-year FFS (**Fig. 3d**), (HR 5.80, p<0.001; Fig. 3c). Among patients with m7-FLIPI data (n=106), the CPC signature improves on m7-FLIPI in predicting poor FFS (C-statistic: 0.834 vs 0.677; **Fig. 3c**). We validated the CPC signature in the independent PRIMA^5^ trial cohort (n=149). High CPC signature enrichment (n=78/149, 52.35%) predicted worse 5-year PFS (**Fig. 3d**).

Together, these results show that the IgM-retaining MBC-like state observed after therapy in mice (cluster-2) has a conserved transcriptional program in human FL that prospectively identifies high-risk patients. The CPC program provides a biologically grounded biomarker for risk stratification and a rationale to target the relapse-founder state.

### An iCPC platform exposes an HDAC dependency

To discover therapies that target the CPC program, we first addressed a limitation of our in-vivo systems, their low throughput for functional and pharmacologic testing. We therefore developed a scalable in-vitro expansion platform that generates CPC-like cells suitable for high-throughput screening. Building on culture paradigms that yield GC-like (iGCB) and memory-like (iMBC) B-cells^38,39^, we optimized conditions to favor the emergence of MBC-like CPCs from the BCK model, establishing an operational iCPC system.

Naïve B-cells from BCK mice were cultured on 40LB feeder expressing CD40L and the pro-survival cytokine BAFF (**Fig. 4a**). Initially exposed to IL-4 followed by IL-21, BCK B-cells adopted a CD95^hi^ CD38^lo^ phenotype, characteristic of GCBs (**Fig. 4b,c**), whereas a tertiary, cytokine-free phase (referred to as the iMBC phase) led to the development of an MBC-like phenotype (CD95^lo^ CD38^hi^) (**Fig. 4a-c**). Compared to iMBCs, iGCB-cells displayed higher proliferation and elevated expression of BCL6 (**Supp. Fig. 2a-d**), a transcription factor essential for GC B-cell identity but not required for MBCs^40,41^.

**Figure 4:**
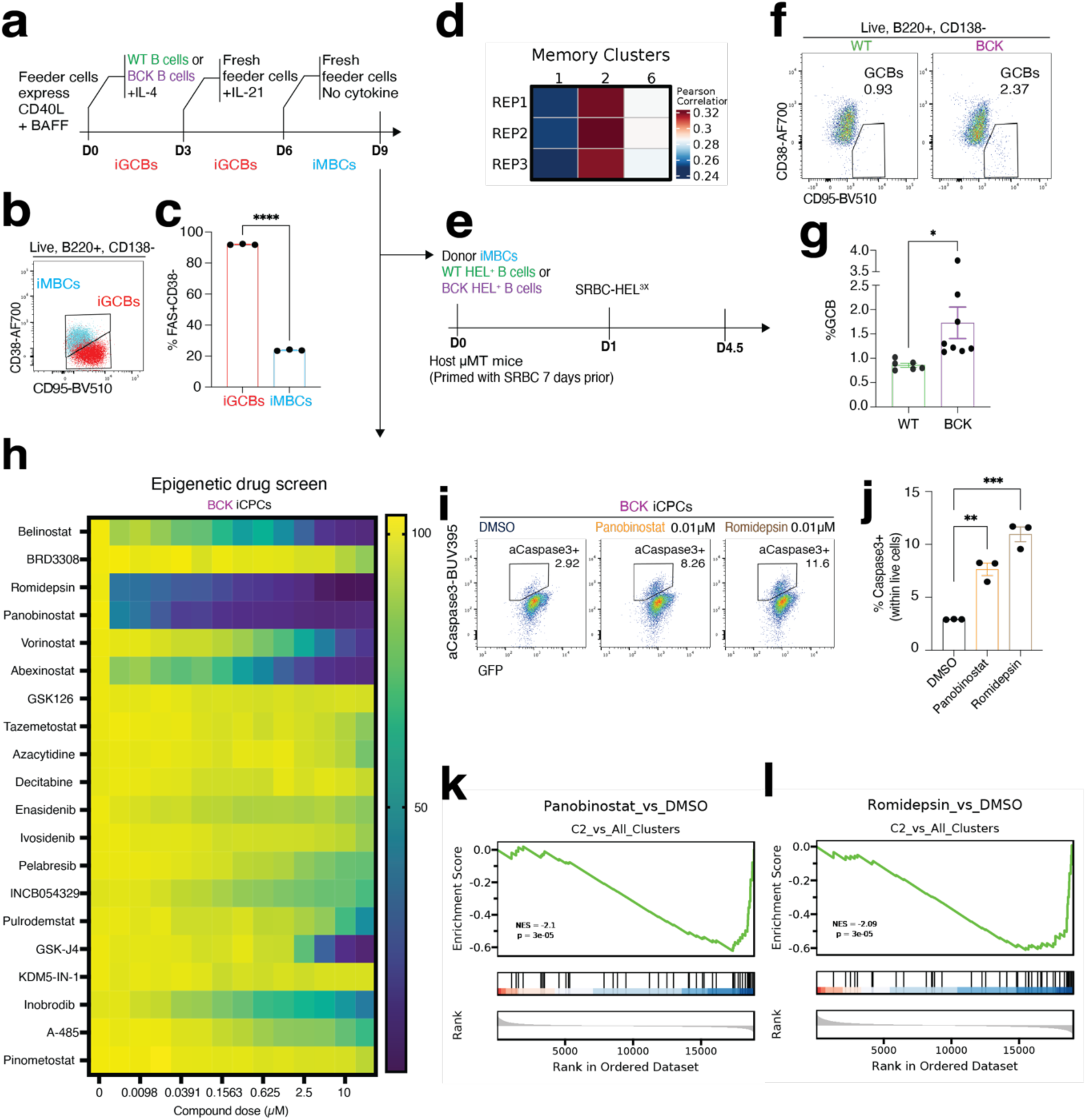
An iCPC platform exposes an HDAC dependency. **(a)** Schematic illustration of the optimized in-vitro germinal center B-cell (iGB) culture system. **(b, c)** Flow cytometry plots and corresponding quantification of iGCB (CD38^lo^FAS^hi^) and iMBC (CD38^hi^FAS^lo^) cells generated *in-vitro*. Data represent mean ± SEM of three technical replicates. **(d)** Heatmap showing transcriptomic similarity between RNA-seq of in-vitro-generated iMBCs and pseudobulk profiles of scRNA-seq-defined MBC clusters 1, 2, and 6. *n* = 3 biological replicates. **(e)** Experimental design for GC re-entry assay: BCK or WT iMBCs were adoptively transferred into SRBC-primed µMT recipient mice, which were subsequently immunized with SRBC-HEL3X. GC re-entry was assessed 3.5 days later. **(f, g)** Flow cytometry gating and quantification showing preferential differentiation of BCK iMBCs into GC B-cells upon antigen recall. Mean ± SEM; WT *n* = 6, BCK *n* = 8. **(h)** Heatmap of iCPC (BCK iMBC) viability after 24-hour treatment with a panel of epigenetic-modifying compounds, measured using the CellTiter-Glo® luminescence assay. Values normalized to DMSO control; mean of three technical replicates shown. **(i, j)** Flow cytometry gating and quantification of cleaved Caspase-3⁺ iCPCs following 24-hour treatment with Romidepsin, Panobinostat, or DMSO. Data shown as mean ± SEM of three biological replicates. **(k, l)** Gene Set Enrichment Analysis (GSEA) of cluster-2 MBC (CPC)-associated genes in HDAC inhibitor (HDACi)-treated vs. DMSO-treated iCPCs. NES = normalized enrichment score; p-values derived from empirical permutation testing; FDR adjusted for gene set size and multiple comparisons. Each dot in (g) and (j) represents an individual mouse. \**P* ≤ 0.05; \*\**P* ≤ 0.01; \*\*\**P* ≤ 0.001; \*\*\*\**P* ≤ 0.0001; ns = not significant (two-tailed Student’s *t*-test).

BCK derived iMBCs exhibited transcriptional similarity to cluster-2 (**Fig. 4d**), the putative CPC population identified in in-vivo models. These cells also demonstrated increased GC re-entry capacity, a hallmark of CPCs^42^. When transferred into μMT recipient mice^43^, BCK iMBCs generated significantly more GC B-cells upon immunization than WT iMBCs, confirming their capacity for GC re-entry (**Fig. 4e-g**), and consistent with a progenitor that repeatedly seeds evolution. These findings support the use of BCK iMBCs (hereafter iCPCs) as functional and transcriptional surrogates for CPCs in in-vitro screens.

Given the early, clonal chromatin-modifier alterations in FL^16^ (**Fig. 1a**), we hypothesized an epigenetic dependency. A focused screen of 20 epigenetic drugs identified HDAC inhibition as toxic to iCPCs (**Fig. 4h**). Both the HDAC1/2-selective inhibitor romidepsin and the pan-HDAC inhibitor panobinostat showed potent single-drug toxicity against iCPCs, with pronounced effects at low nanomolar concentrations (5-10 nM) (**Fig. 4h**). This cytotoxicity was validated by cleaved-Caspase-3 staining, confirming apoptosis induction in iCPCs following romidepsin or panobinostat treatment (**Fig. 4i,j**). To understand the molecular basis of this toxicity, we performed RNA-seq of romidepsin or panobinostat treated iCPCs. GSEA showed that both drugs markedly suppressed the CPC transcriptional program (cluster-2; **Fig. 4k,l**).

Together, these data define a robust, scalable iCPC platform that faithfully models CPC biology and reveals a tractable HDAC dependency. This system enables high-throughput prioritization of CPC-targeted therapies aimed at eradicating the relapse-founding therapy-persistent cells in FL.

### HDAC inhibitors target CPCs in-vivo and in patient-derived organoids

Having identified romidepsin and panobinostat as potent inhibitors of the CPC program, we tested their activity against CPCs in-vivo. In therapeutic model, naïve BCK cells were adoptively transferred into recipients, mice were immunized with HEL, and R-CHOP was given on day 7 (**Fig. 5a**). A single dose of romidepsin, panobinostat, or vehicle was administered on day 11; spleens were analyzed at day 14 (**Fig. 5a**). Both drugs significantly reduced residual BCK cell numbers relative to R-CHOP alone, with panobinostat showing more consistent depletion across animals (**Fig. 5b,c**).

**Figure 5:**
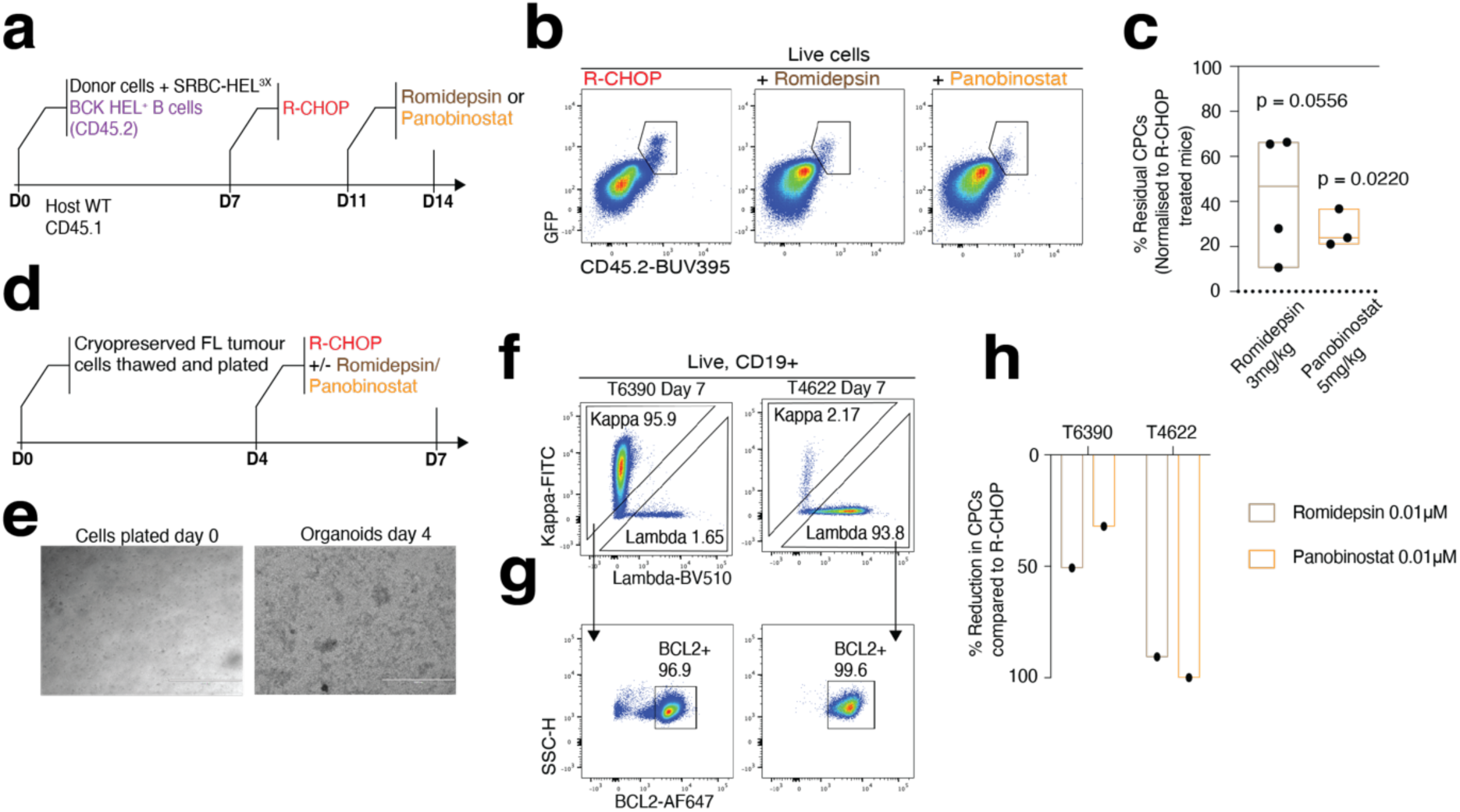
HDAC inhibition targets CPCs in-vivo and patient-derived organoids (a) Experimental design for *in-vivo* assessment of HDAC inhibitor activity. **(b)** Representative flow cytometry plots and gating strategy for quantifying donor-derived BCK cells (GFP⁺ CD45.2⁺) in recipient mice. **(c)** Quantification of residual BCK cells in spleens of mice treated with R-CHOP ± Romidepsin or Panobinostat. Residual cell numbers shown as % of mean BCK cell count in mice treated with R-CHOP alone. Data shown as floating bars (min to max) with line at mean; R-CHOP *n* = 3, Romidepsin *n* = 4, Panobinostat *n* = 3. **(d)** Generation of patient-derived lymphoma organoids (PDLOs): FL biopsies were dissociated, cryopreserved, and subsequently cultured for 4 days prior to treatment with R-CHOP alone or in combination with Romidepsin or Panobinostat. **(e)** Brightfield microscopy images showing the formation of PDLO aggregates after 4 days in culture. Scale bar = 400 μm. **(f, g)** Flow cytometric quantification of viable FL cells in PDLOs on day 7. Lymphoma cells were defined as live, CD19⁺, BCL2⁺ cells with clonally restricted BCR light chains. **(h)** Quantification of treatment effect: reduction in residual FL cell number following Romidepsin or Panobinostat treatment, normalized to R-CHOP–only controls. Each dot in (c) represents an individual mouse. *P*-values calculated using two-tailed Student’s *t*-test.

For human relevance, we evaluated both drugs in patient-derived lymphoma organoids (PDLOs)^44^ established from cryopreserved FL biopsies carrying t(14;18), CREBBP and KMT2D mutations (**Supp. Table 1**). PDLOs received an in-vitro-adapted R-CHOP regimen^45,46^ with concurrent romidepsin, panobinostat, or vehicle control (**Fig. 5d,e**). FL cells within PDLOs were identified as CD19⁺ BCL2⁺ cells with light chain restriction, consistent with the patients’ diagnostic immunophenotype **(Fig. 5f,g**). Both Romidepsin and Panobinostat reduced residual FL cells compared to R-CHOP alone (**Fig. 5h**).

Thus, HDAC inhibition not only kills iCPCs in-vitro but also depletes therapy-persistent residual cells in-vivo and in human organoids, targeting an epigenetic dependency linked to early tumorigenic alterations. These data provide preclinical justification to integrate HDAC inhibitors into front-line, CPC-directed strategies aimed at eliminating the relapse-founder reservoir in FL.

## Discussion

Curing FL will require eradicating the relapse-founding CPC. Progress has been limited by two main barriers: (i) CPC biology was undefined; and (ii) practical tools to prospectively isolate and target this population were lacking. Here, we overcame both with a mouse-first strategy that achieves what is infeasible in patients: immediate post-R-CHOP-like therapy sampling of therapy-persistent cells, prospective isolation of the CPC, and functional testing of its vulnerabilities.

In a genetically faithful early-stage FL model, CPCs map to an IgM⁺ MBC-like state with marked GC re-entry capacity, a behavior consistent with antigenic re-challenge^47^, and the iterative somatic hypermutation-driven clonal evolution that typifies FL^20,35,48,49^. This state is already enriched at diagnosis in patients who experience early relapse and persists after front-line immunochemotherapy within a heterogeneous residual pool. Residual disease is therefore polytypic; survival alone does not equate to relapse potential. Instead, cell identity, captured by the CPC program, better explains outcome. This argues for molecularly defined residual disease to guide risk stratification and intervention.

Although we focused on a t(14;18), CREBBP, KMT2D model because it mirrors the most common early alterations, the CPC program appears to be a convergent genotype-agnostic cell state. CPC-like cells were detectable in a single-cell FL dataset with diverse mutational constellations, and CPC signature enrichment consistently associated with inferior prognosis across large patient cohorts. Clinically, CPC biology offers a biomarker and potential therapeutic target that appears to generalize across mutational backgrounds. Baseline profiling could flag CPC-high patients for risk-adapted intensification or addition of CPC-directed agents, while CPC-signature dynamics may provide a pharmacodynamic marker and a sensitive measure of residual disease to inform monitoring and maintenance. Prospective studies could stratify by CPC burden at diagnosis, pre-specify CPC-signature change as an exploratory endpoint, and evaluate whether CPC-directed add-ons improve outcomes independent of genotype, complementing existing clinicogenetic tools.

Mechanistically, our data suggest state occupancy rather than absolute cell number underlies CPC detection in bulk samples. GC-phenotype cells are eliminated by R-CHOP whereas the CPC program persists, implying a dynamic, treatment-persistent state. CPCs may shuttle between a GC-competent, therapy-sensitive phase and a quiescent, microenvironment-supported phase; genetics define the permissible landscape, while spatial tumor microenvironment cues govern occupancy. Understanding the regulators of these transitions should refine risk, enable dynamic monitoring, and guide timing and choice of CPC-directed therapy.

We find that CPCs coexist with malignant FL cells^13^ and likely occupy multiple anatomical niches, including within secondary lymphoid tissues^50^ and bone marrow^13,14^, with the potential to circulate and persist at non-lymphoid sites^51^. Mapping these niches will clarify spatial heterogeneity, latency, and relapse pathways, and may uncover niche-specific vulnerabilities.

Finally, by operationalizing CPC biology in-vitro (iCPCs) we uncovered a tractable HDAC dependency, with romidepsin and panobinostat ablating iCPCs, depleting R-CHOP persistent cells in-vivo, and in patient-derived organoids. Given that most patients respond to standard therapy, unselected HDAC-inhibitor use is unlikely to yield broad benefit. Our results instead support a precision approach to deploy HDAC inhibitors as CPC-directed add-ons in CPC-high patients identified at diagnosis, and track CPC-signature dynamics as a pharmacodynamic endpoint. Such biomarker-guided trials offer a credible route to eliminating the relapse-founder reservoir, and, ultimately, to durable disease control in FL.

## Methods

### Mice

BCL2 knock-in mice (*Rosa26^LSL^-BCL2.IRES.GFP*) was previously described^22^. To study thereapy-persistent cell development during germinal center (GC) maturation in the context of FL, we integrated these models with the SW_HEL allelic system, allowing precise temporal and antigen-specific control of GC entry following HEL immunization^29^. The following mouse strains were obtained from The Jackson Laboratory (Bar Harbor, ME, USA): C57BL/6J (CD45.2, stock #000664, in house production at Crick Institute), B6.SJL-Ptprc^a^ Pepc^b^/BoyJ (CD45.1, stock #002014, in house production at Crick Institute, *Cd79a-Cre* (stock #020505)^21^, *Crebbp^FL^* (stock #025178)^23^, *Kmt2d^FL^* mice (stock #032152)^24^. *Cd79a-Cre* and *Rosa26^LSL^-YFP* mice were used as controls in relevant experiments. Experiments included both male and female mice, with gender-matching between groups in each experiment. All animals were bred and maintained under specific-pathogen-free conditions at the Francis Crick Institute Biological Research Facility, in accordance with UK Home Office regulations and approved by the Francis Crick Institute Ethical Review Panel.

### Isolation of cells

Single-cell suspensions were prepared from lymphoid tissues, and red blood cells were lysed using ACK lysis buffer. Naïve B-cells were isolated from spleens by negative selection using CD43 (Ly-48) MicroBeads (Miltenyi Biotec), followed by passage of the unlabeled fraction through a MACS LS column (Miltenyi Biotec). This procedure routinely yielded B-cells with >97% purity.

### iGB-cell culture and generation of iMBC cells

Purified B-cells (5 × 10⁵ cells per dish) were cultured in 10-cm tissue culture dishes (BD Falcon) in the presence of mitomycin C–treated 40LB feeder cells (3 × 10⁶ cells per dish; 10 μg/ml, 2 h pre-treatment; Abcam). Cultures were maintained in high-glucose DMEM (Gibco) supplemented with GlutaMAX™, 10% FCS, 5.5 × 10⁻⁵ M 2-mercaptoethanol, 10 mM HEPES, 1 mM sodium pyruvate, 1× MEM non-essential amino acids, 100 U/ml penicillin, and 100 μg/ml streptomycin (Gibco). To generate iGC B-cells, recombinant IL-4 (PeproTech) was added to the primary culture for 3 days. On day 3, cells were re-plated onto a fresh feeder layer and cultured for an additional 3 days with recombinant IL-21 (PeproTech). For tertiary culture, cells were transferred to a new feeder layer without additional cytokines for 3 more days, leading to the generation of iMBC cells for downstream applications and high-throughput drug screening. At each stage, live B-cell numbers were quantified using the trypan blue exclusion method. All cultures were maintained at 37 °C in a humidified atmosphere with 5% CO₂.

### Compound screening

Each compound was initially tested at a starting concentration of 20 μM in a 14-point, 2-fold serial dilution series. Compounds were first prepared in an Echo® Qualified 384-Well Low Dead Volume (LDV) Source Microplate (Labcyte) using the Myra liquid handler (Bio Molecular Systems), with five points of 8-fold dilution in DMSO from a 10 mM stock solution. Volumes of 100 nL, 50 nL, and 25 nL of each dilution were then acoustically dispensed into assay plates (Greiner Bio-One CELLSTAR® 384-well microplates) using an Echo 650T Acoustic Liquid Handler (Beckman Coulter). DMSO backfilling was performed to ensure a uniform final DMSO concentration of 0.2% in a 50 μL assay volume. iMBC cells were harvested from 40LB feeder layers using Cell Dissociation Buffer (Gibco) and resuspended in B-cell media supplemented with recombinant mouse BAFF (200 ng/ml; Bio-Techne) and recombinant mouse CD40L (1 μg/ml; Bio-Techne), at a final density of 2 × 10⁵ cells/mL. A total of 10,000 cells in 50 μL were seeded into each well using a ViaFill dispenser (Integra). After 24 hours of incubation at 37 °C with 5% CO₂, cell viability was assessed by measuring intracellular ATP levels using the CellTiter-Glo® assay (Promega). Following the addition of 40 μL of CellTiter-Glo reagent, plates were incubated at room temperature for 20 minutes before luminescence was read using the EnSight plate reader (Revvity). Viability was expressed as a percentage relative to DMSO-treated control wells.

### Immunization, treatments, and adoptive transfers

As previously described^29^, splenic cells from donor mice were incubated with CD43 (Ly-48) MicroBeads (Miltenyi Biotec; 130-049-801) and enriched for B-cells by negative selection using MACS LS columns (Miltenyi Biotec). The resulting CD19⁺B220⁺ population was routinely >97% pure. To quantify the frequency of HEL-specific B-cells, 0.5 × 10⁶ cells were incubated with 200 ng/ml HEL on ice for 20 minutes, washed twice with FACS buffer, and stained with HyHEL9-Alexa Fluor® 647. HEL-binding B-cells were quantified by flow cytometry. A total of 1-2 × 10⁶ HEL-specific B-cells were transferred intravenously into CD45.1⁺ B6.SJL recipient mice. On the same day, mice were immunized intravenously with 2 × 10⁸ SRBCs conjugated to HEL3X. HEL3X protein was produced in Chinese hamster ovary (CHO) cells expressing His-tagged HEL3X. The protein was purified from culture supernatant using a HisTrap FF column (Cytiva) on an ÄKTA Pure chromatography system (Cytiva). Eluted fractions were concentrated to 0.5 ml using a Vivaspin 15 concentrator (10 kDa cutoff), then subjected to size-exclusion chromatography on a Superdex 200 10/300 column (Cytiva). HEL3X eluted as a single symmetrical peak, was snap-frozen at 1 mg/ml, and stored at-80 °C until use.

### R-CHOP treatment

Ultra-LEAF™ Purified anti-mouse CD20 antibody (BioLegend) intravenously, Cyclophosphamide intravenously, Doxorubicin intravenously, Vincristine intravenously and Prednisolone oral gavage daily for 5 days, dosing of chemotherapy was based on previous work^52^. Panobinostat (BPS Bioscience) for intraperitoneal injection was dissolved in 5% DMSO (Sigma), 40% PEG300 (MedchemExpress), 5% Tween 80 (Sigma-Aldrich), 50% ddH_2_0. Romidepsin (Active Motif) for intraperitoneal injection was dissolved in 10% DMSO, 90% saline.

The percentage of B-cells in S-phase was assessed using an EdU incorporation assay (Invitrogen). Mice were administered 1 mg EdU dissolved in PBS via intraperitoneal injection 1 hour prior to euthanasia. Following surface marker staining, splenocytes were fixed, permeabilized, and subjected to a click chemistry reaction to fluorescently label incorporated EdU using the Click-iT™ Plus EdU Flow Cytometry Assay Kit with Pacific Blue detection (ThermoFisher; C10636), according to the manufacturer’s instructions.

### Flow cytometry analysis and cell sorting

Single-cell suspensions of mononuclear splenocytes were processed for flow cytometry as follows: cells were first incubated with anti-mouse CD16/32 antibody (BD Biosciences; 553142; 1:200 dilution) in FACS buffer for Fc receptor blockade (10 minutes at 4 °C). Dead cells were excluded using Zombie NIR viability dye (BioLegend; 423106) incubated in FACS buffer for 20 minutes at 4 °C. Surface staining was performed using a panel of fluorochrome-conjugated anti-mouse antibodies diluted in FACS buffer and incubated for 30 minutes at 4 °C. To identify HEL-binding B-cells, cells were stained with HEL protein (50 ng/mL) followed by HyHEL9-Alexa Fluor® 647 detection. For intracellular staining, cells were fixed and permeabilized using the True-Nuclear™ Transcription Factor Buffer Set (BioLegend), followed by overnight antibody incubation at 4 °C. Flow cytometry acquisition and analysis were performed using an LSR Fortessa or FACSAria (BD Biosciences). Data were analyzed using FlowJo v10.8.1. For fluorescence-activated cell sorting (FACS), antibody-labeled cells were sorted on a BD FACSAria II equipped with five lasers (355, 405, 488, 561, 640 nm).

The anti-mouse antibodies were diluted as follows: BUV395 anti-CD45.2 (BD 564616, Clone 104, 1:200), BV510 anti-CD95 (BD 563646, Clone Jo2, 1:200), R718 anti-CD95 (BD 752226, Clone Jo2, 1:200), PE-Cy7 anti-CD95 (BD 557653, Clone Jo2, 1:200), AF700 anti-CD38 (Invitrogen 56-0381-82, Clone 90, 1:200), BV650 anti-CD38 (BD 740489, Clone 90, 1:200), BV605 anti-IgD (BD 405727, Clone 11-26c.2a, 1:200), BV421 anti-IgD (BD 405725, Clone 11-26c.2a, 1:200), AF594 anti-IgD (BD 405740, Clone 11-26c.2a, 1:200), PerCP-EF710 anti-IgM (Invitrogen 46-5790-82, Clone II/41, 1:200), BUV805 anti-IgM (BD 749307, Clone II/41, 1:200), Biotin anti-Active Caspase-3 (BD 550557, Clone C92-605, 1:10), PE-Cy7 anti-BCL6 (BD 563582, Clone K112-91, 1 in 1:20), PE anti-Active Caspase-3 (BD 570183, Clone C92-605, 1:20), PerCP-EF710 anti-CXCR4 (Invitrogen 46-9991-82, Clone 2B11, 1:200), BV510 anti-B220 (BioLegend 103248, Clone RA3-6B2, 1:200), BUV737 anti-B220 (BD 612838, Clone RA3-6B2, 1:200), BV605 anti-CD86 (BioLegend 105037, Clone GL-1, 1:200), BV786 anti-CD138 (BD 740880, Clone 281-2, 1:200), BUV737 anti-Kappa LC (BD 748499, Clone 187.1, 1:200), AF700 anti-Kappa LC (BioLegend 409508, Clone RMK-45, 1:200). The anti-human antibodies were diluted as follows: FITC anti-Kappa LC (BioLegend 316506, Clone MHK-49, 1:50), BV510 anti-Lambda LC (BD 570934, Clone 1-155-2, 1:50), AF647 anti-BCL2 (BioLegend 658706, Clone 100, 1:20), BV650 anti-CD19 (BioLegend 302238, Clone HIB19, 1:100).

For FACS sorting, cells were incubated with antibodies and sorted on BD Aria II sorter with 5 lasers (355, 405, 488, 561, 640). Zombie NIR (Biolegend 423106) was used to exclude dead cells in all experiments.

### Histopathology and immunohistochemistry

Tissue was fixed for 24 h in 10% Neutral Buffered Formalin (NBF) before processing to wax using a Tissue-Tek VIP 6 AI processor. 3 μm tissue sections baked for 1h at 60°C prior to immunohistochemical staining using on a Roche Ventana Discovery Ultra autostainer for rabbit anti-CD3 (Abcam, ab134096) or rabbit anti-CD19 (Abcam, ab245235), or rabbit anti-BCL6 (Cell Signaling Technology, 5650S) with DISCOVERY OmniMap anti-Rb HRP (RUO) (Roche, 760-4311) and detected with DISCOVERY ChromoMap DAB Kit (RUO) (Roche, 760-159). DISCOVERY CC1 solution (Roche, 950-500) was used to retrieve the targets. Slides were counterstained with haematoxylin using a Tissue-Tek Prisma automated slide stainer. Slides were coverslipped using Tissue-Tek Glas g2 Automated Glass Coverslipper.

### Single-cell processing of B-cells from BCK mice

Raw sequencing reads from 10x Genomics were aligned to the Ensembl 98 genome and quantified using the *Cellranger multi (v7.1.0)* pipeline. Count matrices from placebo-treated, RCHOP-treated and tumor samples were subsequently processed together with *Scanpy (v1.9.4)* in *Python (3.10.12)*, where genes with low capture rate (< 5 cells) were removed. Sample demultiplexing was conducted exclusively on the R-CHOP-treated runs to identify single cells from individual mice with the ‘*MULTIseqDemux()’* function. Cells flagged as doublets or negatives by demultiplexing were subsequently excluded from the count matrix. Additional stringent thresholds were applied to remove droplets with <800 *nFeature_RNA*, and >5 median absolute deviation (MAD) in *nFeature_RNA, nCount_RNA,* and *percent.mt*.

To ensure the quality of B-cells in our analysis, droplets contaminated with high hemoglobin reads (*percent.hb* >0) and TCR reads (*percent.tcr* >0.05%) were removed. Cells were normalized with the shifted logarithm method, and 3000 highly variable features (HVFs) were selected for principal component analysis (PCA). Preliminary clustering was conducted using the first 20 PCs and Leiden algorithm at varying resolutions for each group (placebo=0.5, RCHOP=0.3, tumor=0.5), where clusters corresponding to T/NK-cells (*Cd3d, Nkg7*), myeloid cells (*Lyz2, Itgam*), and plasma cells (*Irf4, Prdm1*) were removed from all treatment groups. After quality control, 58,497 B-cells remain across all treatments.

### Integration & analysis of B-cells from BCK mice

High quality B-cells were normalized using the analytical Pearson residual method with 3000 HVFs. By performing PCA and Harmony integration across samples from treated and tumor B-cells, we used the first 20 PCs to calculate cell-cell nearest neighbors and conducted Leiden clustering with a resolution of 0.6. Single cells were projected onto a 2D space using uniform manifold approximation and projection (UMAP) for visualization. B-cell clusters were manually annotated based on the selected marker expression: activated (*Dusp2, Cd69, Fos*), memory (*Hhex, Cd38, Klf2*), germinal center light zone (*Il4i1, Cd40, Fcer2a*) and dark zone *(Mef2b, Aicda, Bcl6*).

### Copy number variation inference of BCK B-cells

Copy number variations (CNVs) in B-cells from BCK mice were analyzed using the *inferCNV* algorithm *(v1.21.0)* with the following parameters: *cutoff = 0.1, denoise = TRUE, and HMM_type = “i6”*. B-cells from WT mice served as the normal-cell reference to establish baseline expression levels. To calculate the CNV correlation score, we averaged the CNV profiles from tumor samples and computed the Pearson correlation coefficient for every single cell.

### Single-cell differential expression analysis

Differential gene expression per cluster was calculated in BCK tumor samples with the *‘FindAllMarkers()’* function from *Seurat (v5.0.1)* using a zero-inflated generalized linear model *(MAST, v1.26.0)* method, implemented in *R (v4.3.2)*. Genes were p-adjusted by the Bonferroni method and considered significantly upregulated with *p_val_adj of < 0.001, pct.1 > 0.1* and *avg_log2FC > 0.5*.

### Processing and reference mapping of FL scRNA-seq dataset

FL patient-derived B-cells from Radtke et al. were normalized per sample using the SCTransform method using the following arguments: *vst.flavor = “v2”*, *variable.features.n = 3000*. RPCA-based batch correction was performed on a sample level with *FindIntegrationAnchors()* and *IntegrateData()* from the *Seurat V4* pipeline, with the *number of integration features* and *k.weight* set to *3000* and *40* respectively.

To identify homologous cell types across species, Seurat’s *MapQuery()* function with *dims = 1:20* and *method = “rpca”* was employed to classify patient B-cell subsets and project them onto the integrated murine UMAP space.

### RNAseq of B-cells treated with HDAC inhibitors

Fasta files for RNAseq were aligned to the GRCm39 reference genome (Ensembl 98) using the nf*-core* pipelines *(v3.13.2)*, with transcripts aligned and quantified using the STAR-Salmon method. Variance-stabilizing transformation (VST) was applied to the transcripts for data normalization. For each gene, the mean expression across all samples was calculated, and a threshold corresponding to the 5th percentile *(quantile = 0.05)* of this distribution was used to filter out lowly expressed genes. Differential expression analysis was conducted using *DESeq2 (v1.40.2)* with default settings, followed by fold change shrinkage using *ashr (v2.2-63)*. Genes were considered differentially expressed with *log2FoldChange >|0.5|* and *padj < 0.05*.

### Gene set enrichment analysis

Differentially expressed genes from all datasets were pre-ranked based on log2 fold change expression. Cluster signatures, as well as signatures from *MSigDB HALLMARK, KEGG, GOBP and REACTOME* collections were collated to perform gene set enrichment analysis using the *GSEA()* function from the *clusterProfiler* package *(v4.8.2)*, with parameters: *nperm = 1000, min.overlap = 5*. Pathways are considered significant when *q-value < 0.05*.

### FL Organoids

Patients undergoing treatment and care at Barts Health NHS Trust were consented for an observational study of blood and tissue-based biomarkers approved by the local IRB (Ethics ref no: 21/EE/0123). All patients were provided with written informed consent per protocol. Tumor biopsy samples were obtained as indicated by standard of clinical care at diagnosis from individuals with diagnosed or suspected lymphoma. Treatment decisions were made at the discretion of the treating physician in accordance with institutional standards. Patients included in this study were enrolled between 2012 and 2020. A summary of patient age, sex, disease type, and clinical variables is provided in Supplementary table S1. FL organoids were generated as previously described^44^. Briefly, surgically excised lymph nodes containing viable lymphoma cells were collected as fresh tissue in saline or PBS at diagnosis. Samples were mechanically dissociated in DPBS over ice utilizing a sterile pestle and stainless-steel cell dissociation sieve into single-cell suspensions, resuspended in freezing media (10% DMSO, 50% FBS, 40% RPMI), and stored in liquid nitrogen until preparation as immune organoids. To generate organoids, cryopreserved cells were thawed, and excess DNA was removed with benzonase treatment (1:10,000 in media while thawing cells; Sigma). Cells were washed with RPMI+10% FBS and plated at a final density of 7.5 x10^6^/ml (100μl final volume; 7.5×10^5^ live cells/well) in flat 96-well ultra-low attachment plates (Corning). Organoid media was composed of RPMI1640 with glutamax, 10% FBS, 1x nonessential amino acids, 1x sodium pyruvate, 1x penicillin–streptomycin, 1x Insulin-Transferrin-Selenium (ITS - G) supplement (Gibco). Cultures were incubated at 37°C, 5% CO2 with humidity and media was replenished every other day by topping up with 30 μl of fresh organoid media. On day 4 of culture, organoids were treated with an in-vitro adapted R-CHOP based on clinically achievable concentrations of doxorubicin, and Rituximab administered in the R-CHOP regimen^45,46^.

### Statistical analysis

Statistical methods used for p value calculation were specified in the corresponding figure legends. Statistical analyses were conducted using either GraphPad Prism 9 or the R statistical language scripts and packages specified. Data were judged to be statistically significant when p or adjusted p < 0.05 unless otherwise noted. All experiments were successfully performed with at least three biological replicates unless otherwise stated in the figure legend. No statistical method was used to predetermine sample size. Occasionally, specific samples were excluded from analyses because of technical flaws in processing or data acquisition. Mice were randomised into treatment groups by age-and sex-matching before tumor inoculation. The data were analyzed by groups and the investigators were blinded to group allocation during experiments. For all data analysed with two-tailed Student’s *t*-test, normality was checked using Kolmogorov-Smirnov test or Shapiro-Wilk test where small sample size prevented use of Kolmogorov-Smirnov test. For non-normally distributed data a two-tailed Mann Whitney test was used. Sample sizes were chosen based on prior studies and pilot experiments.

## Acknowledgements

We thank the members of the Immunity and Cancer laboratory [Francis Crick Institute (FCI), London, UK] for critical discussions and comments. We thank John Riches [CRUK Barts Cancer Institute, QMUL] for critical discussions and comments. We thank the FCI scientific platforms (Biological Resource Facility, Flow Cytometry, Histopathology) for expert advice and technical support. We acknowledge the Genomics Science Technology Platform at The Francis Crick Institute, for their contribution to the single-cell capture experiment, library construction, and sequencing, particularly Hubert Slawinski, Deb Jackson, Sam Jones and Marg Crawford. Schematics were created with <u>BioRender.com</u>

## Author contributions

Conceptualization: O.A., M.S.H., J.F., J.O., L.Z., and D.P.C.

Methodology: O.A., M.S.H., L.Z., O.S., B.M.

Investigation: O.A., M.S.H., L.Z., O.S., B.M., B.C., C.E.

Resources: B.S., S.H., G.S., M.H., H.C.R., J.O.

Visualization: O.A., M.S.H., L.Z.

Funding acquisition: H.C.R., J.F., J.O., O.A., D.P.C.

Supervision: L.Z., and D.P.C.

Writing-original draft: O.A., M.H., L.Z., and D.P.C.

Writing-review and editing: All authors.

## Supplementary Figures

**Supp. Figure 1.**
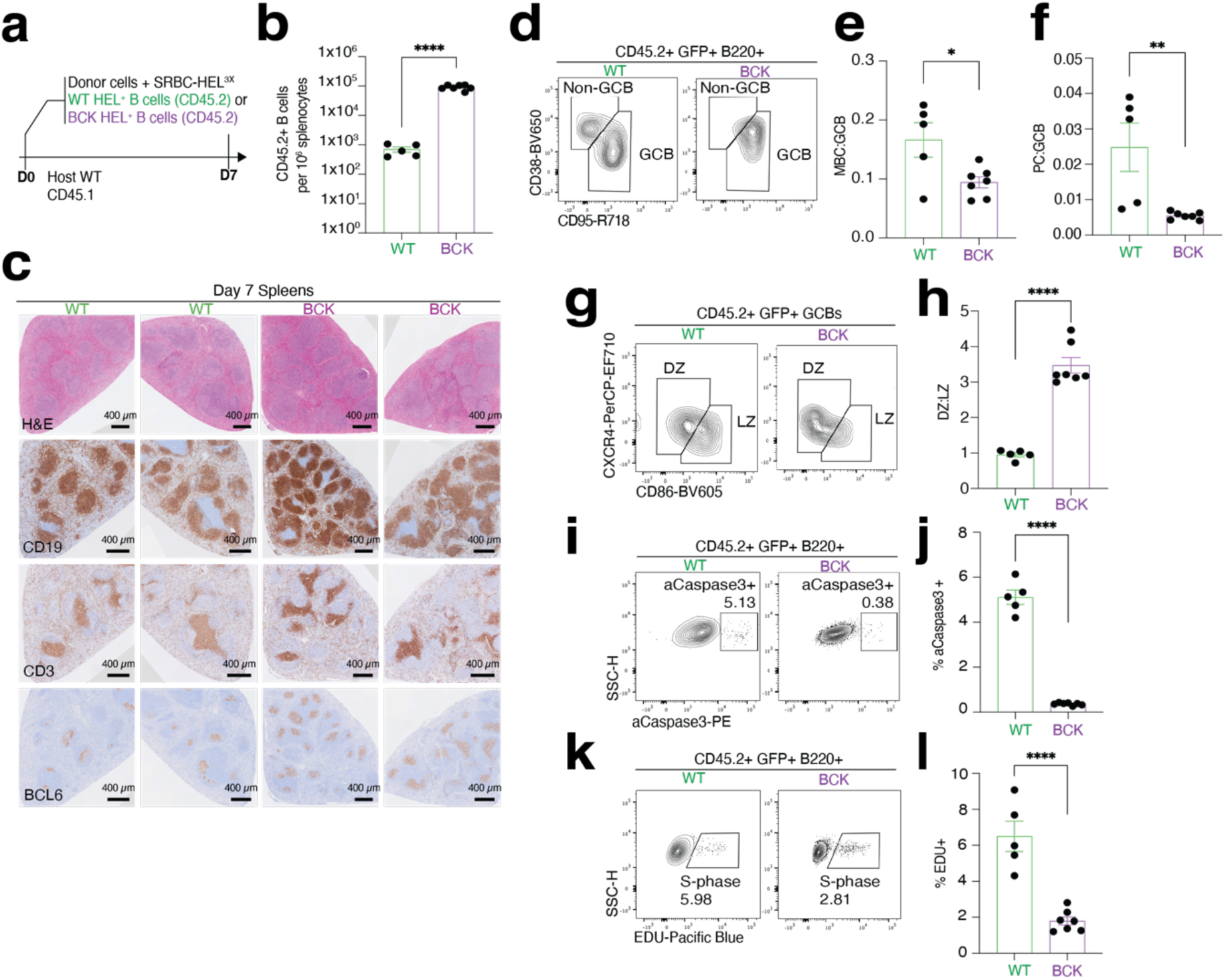
GC hyperplasia, reduced apoptosis and differentiation in BCK. (a) Experimental design for adoptive transfer and immunization.= **(b)** Quantification of donor-derived (CD45.2⁺) B-cells per 10⁶ splenocytes in WT and BCK recipients. Mean ± SEM; WT *n* = 5, BCK *n* = 7. **(c)** Representative immunohistochemistry (IHC) images of spleen sections showing GC expansion. Scale bar = 400 μm. **(d)** Flow cytometry gating strategy for identifying GC B-cells (GCB) and non-GCB populations. **(e, f)** Ratios of memory B-cells (MBC) to GC B-cells and plasma cells (PC) to GC B-cells, assessed by FACS on day 7 post-immunization. Mean ± SEM; WT *n* = 5, BCK *n* = 7. **(g, h)** Gating strategy and quantification of dark zone to light zone (DZ:LZ) ratio within GCB populations. **(i, j)** Proportion of apoptotic B-cells, as defined by active caspase 3. **(k, l)** Proportion of B-cells in S-phase, indicating proliferative activity. Each dot represents an individual mouse. \**P* ≤ 0.05; \*\**P* ≤ 0.01; \*\*\**P* ≤ 0.001; \*\*\*\**P* ≤ 0.0001 (two-tailed Student’s *t*-test); ns, not significant.

**Supp. Figure 2:**
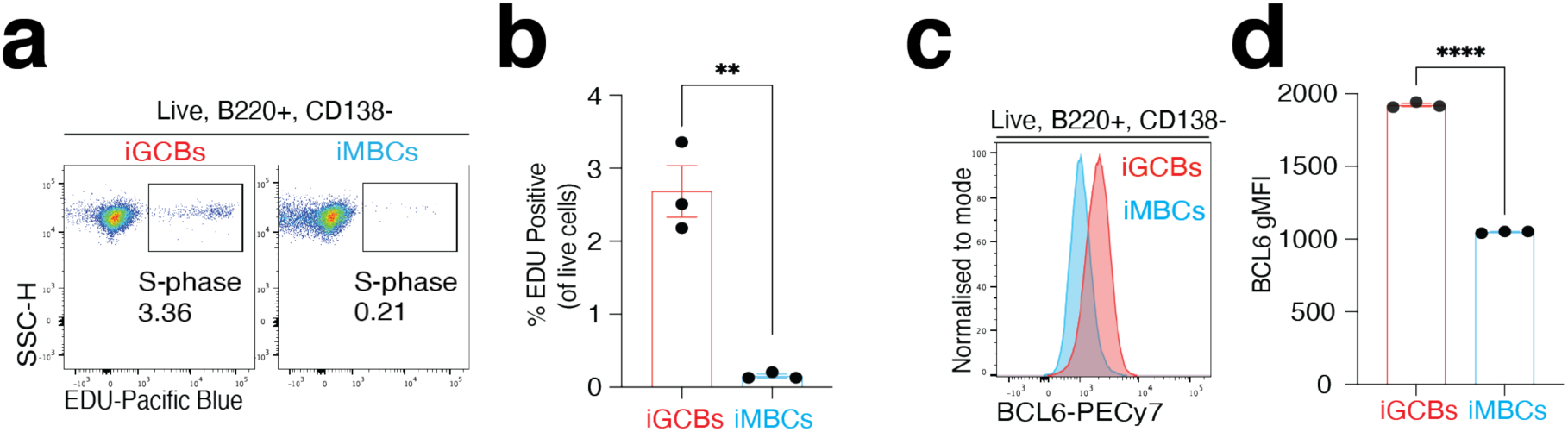
Phenotype of BCK iCPCs. (a, **b)** Representative flow cytometry plots and quantification of the proportion of in-vitro–generated GC B-cells (iGCB) and memory B-cells (iMBC) in S-phase, indicating proliferative status. Mean ± SEM; *n* = 3. **(c, d)** BCL6 expression analysis in iGCB and iMBC cells, presented as geometric mean fluorescence intensity (gMFI). Data represent mean ± SEM from three technical replicates. Each dot in (b) represents an individual mouse. \**P* ≤ 0.05; \*\**P* ≤ 0.01; \*\*\**P* ≤ 0.001; \*\*\*\**P* ≤ 0.0001 (two-tailed Student’s *t*-test); ns = not significant.

---

**Supp. Table 1:** Clinical characteristics of cohort used to generate organoids.

| Patient ID | Mutation profile | Gender | Disease state | Type | Age at diagnosis | Stage at diagnosis | FLIPI at diagnosis | Previous treatment |
| --- | --- | --- | --- | --- | --- | --- | --- | --- |
| T6390 | KMT2D (G5428D and A1390fs), CREBBP (Y1412N), RRAGC | Female | Pre-treatment | FL non-transformed | 40 | IV | 2 | Expectant management, R-CHOP + R-maintenance |
| T4622 | KMT2D, CREBBP, EP300 | Male | Diagnosis | FL transformed | 48 | IV | 4 | R-CHOP + R-maintenance, R-ICE, mini-BEAM, R-GEM-P, Allo-SCT |

